# Single-cell transcriptome profiling simulation reveals the impact of sequencing parameters and algorithms on clustering

**DOI:** 10.1101/2021.03.16.435626

**Authors:** Yunhe Liu, Bisheng Shi, Aoshen Wu, Xueqing Peng, Zhenghong Yuan, Gang Liu, Lei Liu

## Abstract

Despite of scRNA-seq analytic algorithms developed, their performance for cell clustering cannot be quantified due to the unknown “true” clusters. Referencing the transcriptomic heterogeneity of cell clusters, a “true” mRNA number matrix of cell individuals was defined as ground truth. Based on the matrix and real data generation procedure, a simulation program (SSCRNA) for raw data was developed. Subsequently, the consistence between simulated data and real data was evaluated. Furthermore, the impact of sequencing depth, and algorithms for analyses on cluster accuracy was quantified. As a result, the simulation result is highly consistent with that of the real data. It is found that mis-classification rate can be attributed to multiple reasons on current scRNA platforms, and clustering accuracy is not only sensitive to sequencing depth increasement, but can also be reflected by the position of the cluster on TSNE plot. Among the clustering algorithms, Gaussian normalization method is more appropriate for current workflows. In the clustering algorithms, k-means&louvain clustering method performs better in dimension reduced data than full data, while k-means clustering method is stable under both situations. In conclusion, the scRNA simulation algorithm developed restores the real data generation process, discovered impact of parameters on mis-clustering, compared the normalization/clustering algorithms and provided novel insight into scRNA analyses.

## INTRODUCTION

Single-cell RNA sequencing technology has developed rapidly in recent years, and has gradually become preferred sequencing technology for researchers in the fields including histological variation^[1–3]^ and tumor immune microenvironment^[4, 5]^. However, there are still some shortcomings in current analysis workflow. The sequencing depth has now reached over 1 billion reads in one scRNA-seq sample of recent studies^[3]^, while after assigning to about 20,000 cells captured, the average reads allocated to a single cell (50,000) is insufficient for transcriptome profiling^[6]^. The impact of low coverage on the cell classification is difficult to quantification. Thus, quantifying the sequencing depth and clustering accuracy is necessary for scRNA analytic procedure.

In the past years, multiple scRNA-seq analytic algorithms have been proposed, according to reasonable assumptions and models, including, Biscuit, K-means&louvain in clustering, and MNN, CCA in batch effect removing^[7–9]^. However, compared with other NGS based technologies, few articles used exactly the same analysis workflow or parameter^[1–5, 10–12]^. For the same data, using different algorithmic may lead to quite different results. And in the absence of ground truth, the researchers might choose algorithms subjectively.

However, the relationship between clustering accuracy and depth/algorithms cannot be effectively estimated due to the unknown “true” clusters. Simulation is a frequently used option. With pre-defined ground truth and parameters, the influence of parameters can be quantified without systematic errors^[13]^. Currently, several simulation programs for single cell data have been released, including SPsimSeq, Splatter, SPARSim and SymSim^[14–17]^. These algorithms are all hypothesis-driven, instead of data-proposed models^[18, 19]^. The drawback for the algorithms is that consistence between real data and simulation data is not satisfactory. For single cells with more diverse expression profiles among different cells^[20]^, it is difficult to use a single mathematical model to fit or formulate a ground truth.

To address the problems, an SSCRNA program (https://github.com/liuyunho/SSCRNA-v1.0) was developed, following pre-defined ground truth, to simulate scRNA-seq data (fastq data). The SSCRNA program mimics the actual sequencing process including the sequencing library building and sequencing process^[21]^, which enables flexibility to adjust the parameters that may be introduced in each step of sequencing and like in real process, a constructed sequencing library can be used for several sequencing with different paraments. The reliability of SSCRNA program was verified by comparing the analysis results of the real data with simulation data. Using this tool, the impact of sequencing depth on clustering accuracy was quantified, and the performance of current analysis procedures was also evaluated.

## RESULTS

### SSCRNA - a simulation program to generate scRNA-seq data

With the development of scRNA-seq technology, a number of available sequencing platforms have emerged, including SMART-seq2, CELL-seq and Drop-seq^[22–24]^, all of which consist of three sections: cell isolation and capture, library building, and sequencing (Figure 1A 1–3).

**Figure 1.**
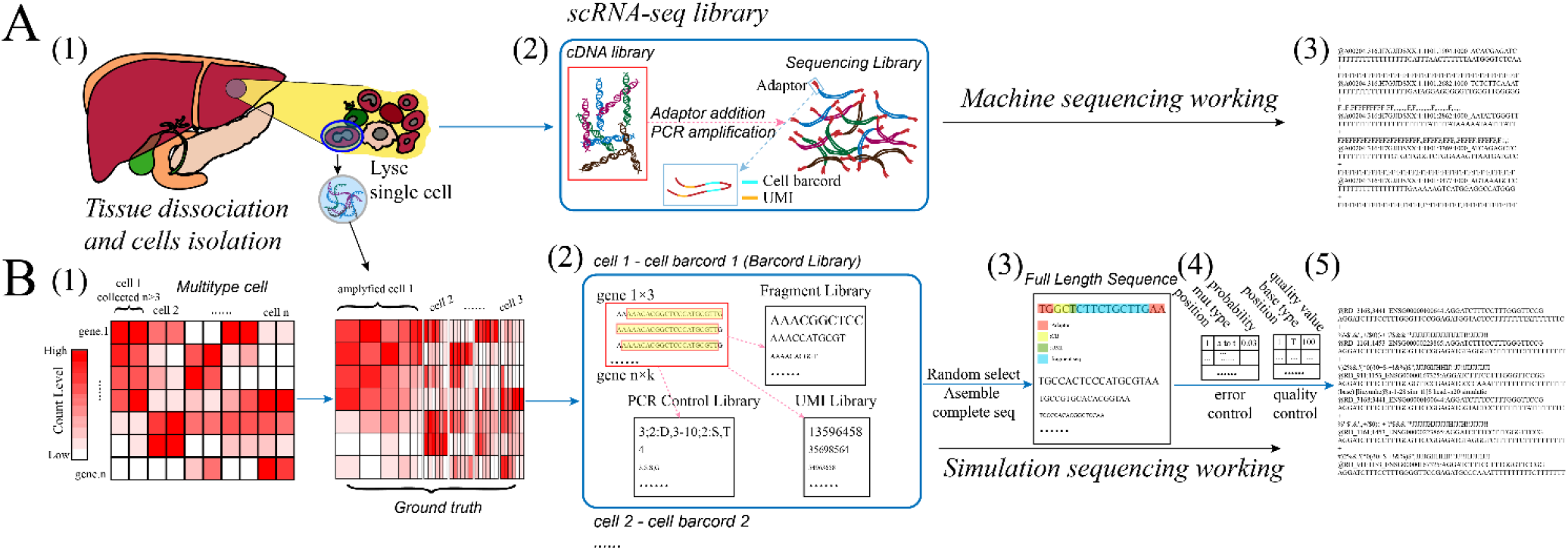
(A) Diagram of the actual scRNA-seq process (1. Tissue dissociation, cell isolation and capture; 2. ScRNA-seq library building; 3. Data generated by sequencing). (B) Diagram of the overall process of the SSCRNA program (1. Pre-defined ground truth. (by amplifying the expression data of collected enriched cells);2. Simulation sequencing library; 3. Full length sequence; 4. Error and quality control files; 5. Data generated by SSCRNA program).

A pre-defined ground truth was constructed in the first (Figure 1B 1, as a reference for the expression profile of captured single cells, in other words, an expression matrix was generated containing the numbers of mRNAs in each cell). Then, the SSCRNA program mimics the actual sequencing process, utilizes the ground truth to perform the simulation of remaining sections (Figure 1B 2–5; see Materials and Methods). In library building, the SSCRNA implements tag-based quantification method^[25]^, assigning the corresponding fragment sequences, UMI tags and cell barcodes to sub-library files and using PCR control sub-library file to record the number and mutation of the sequences after amplification simulation (Figure 1B 2). In subsequent sequencing simulation, the program randomly selects sequences from the 4 sub-library files, and assembles the corresponding fragments into full length sequence (Figure 1B 3). The reading error and basis quality in read generation process are controlled by error control file and quality control file respectively (Figure 1B 4). After the steps mentioned above, the simulation scRNA-seq raw data (fastq format) are generated (Figure 1B 5).

### The comparison of the real data with the simulated data

The analysis result of a part of real dataset (DA3, Table S1) was used as ground truth for simulating, which means that the simulation is based on the real condition for simulation evaluation. The aforementioned SSCRNA program was employed to perform the simulation process to generate the simulation data (Table S2, see Materials and Methods). The consistence between simulated and real data was evaluated. In the cell barcode estimation step, the knee point of the cell barcode-count curve^[26]^ is a commonly used indicator for true/noise distinguishing (Figure 2A). The profile of barcode-count between real/simulated data is similar. The corrective effect^[27, 28]^ of the UMI addition in simulation data was verified. The results (Figure 2B) shown that as the number of genes gradually increasing, the difference between the mapping count and the real count gradually increases. Clustering result between simulated and real data was compared, as shown in Figure 2C, the cell cluster distribution profile between real and simulated data is similar. Cluster-specific genes in real and simulated data is identified, and the overlapped genes were shown in Figure 2D. As expected, the corresponding cluster-specific genes could be re-detected in each simulated cell cluster. Together, these results indicate that the SSCRNA program reproduces the real data.

**Figure 2.**
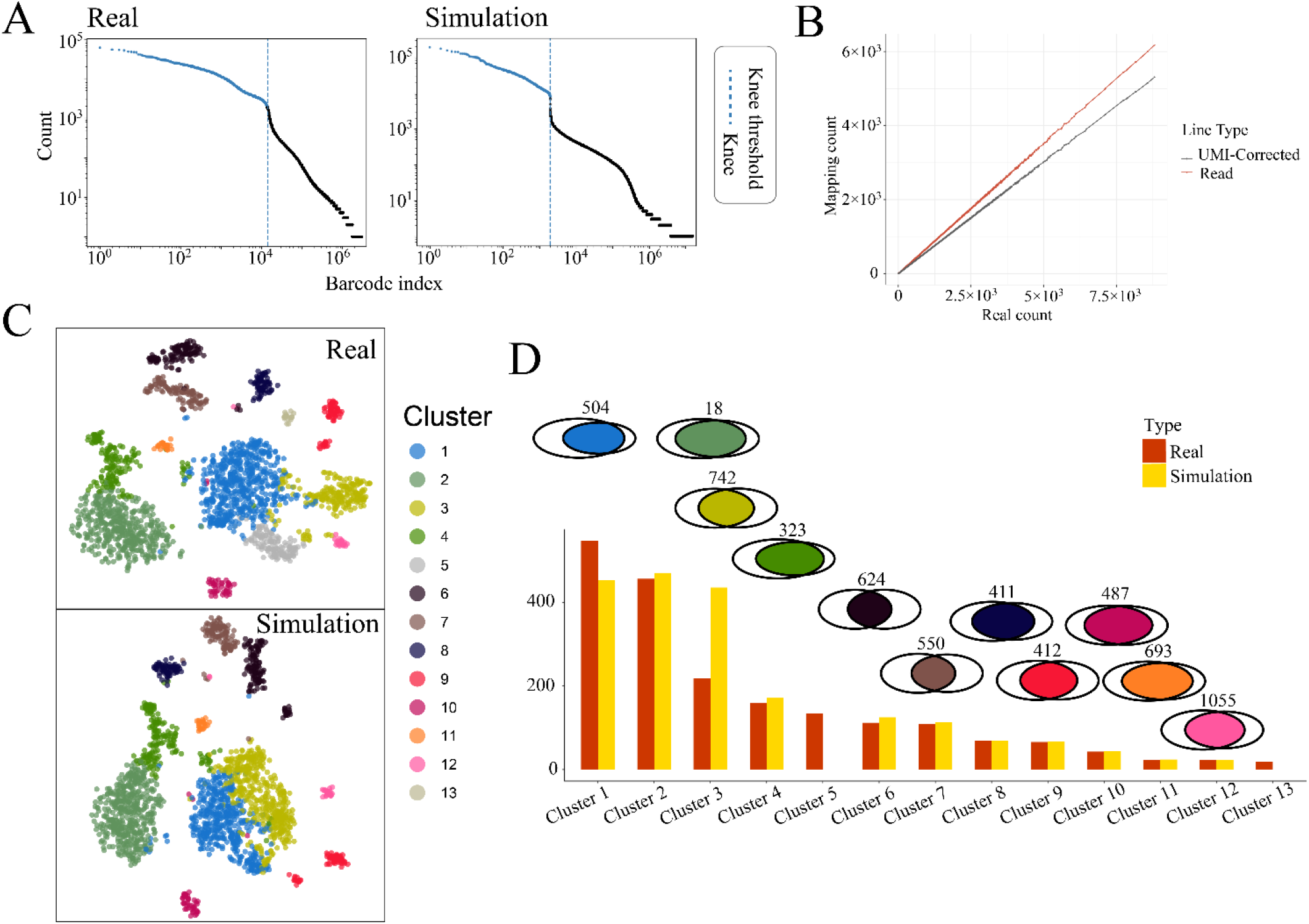
(A) Diagram of the cell barcode-count curve (left panel: real data; right panel: simulation data). (B) Diagram of mapping count-real count plot with or without UMI correction. (C) TSNE plot (upper panel: real data; lower panel: simulation data). (D) Comparison between corresponding cluster of real and simulation data. (Boxplot: cell number; Venn diagram: interaction of specific genes, and the colour of the intersection part is consistent with the colour of the cluster in the TSNE plot)

### Impact of sequence depth on classification accuracy

A collected dataset containing 11 major-categories and 42 sub-categories of immune cell individuals (Table S3; At least three samples per sub-category) was used. Correlations between sub-categories individuals were calculated (Figure 3A; left panel: using whole genes; right panel: using 530 hemocyte-specific genes^[29]^; see Materials and Methods). Intra-category samples have significantly higher correlations than extra-category, regardless of using all features or optimized features. The ground truth is obtained by amplifying each sub-category to 50 individuals each (see Materials and Methods). Gradient sequencing depth was set for simulation data generation (Table S4; 4 main gradients; 8 sub-gradients).

**Figure 3.**
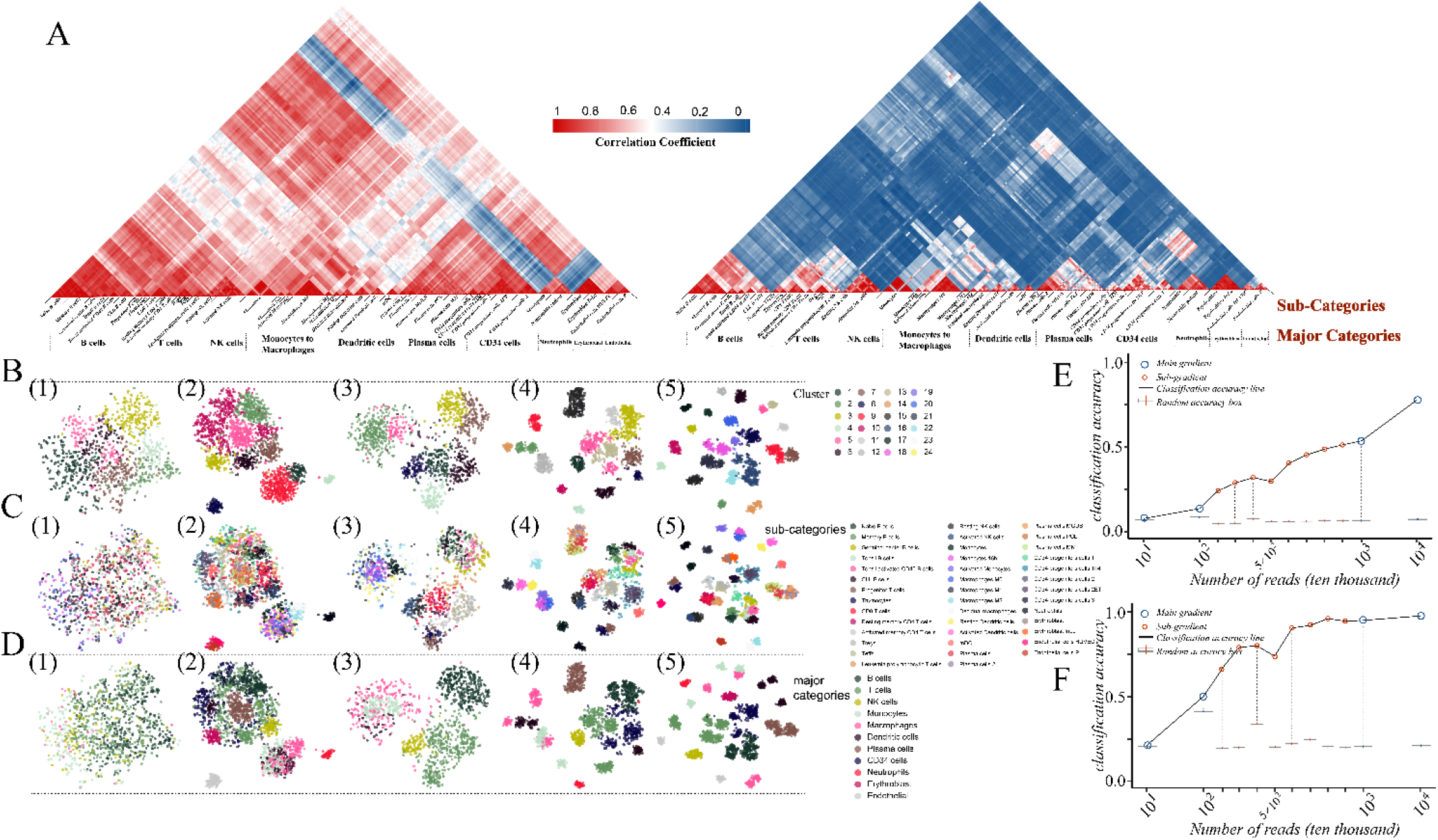
(A) The correlation heatmap of combined data. (1. correlation calculated by whole genes; 2. by 530 hemocyte-specific genes) (B) TSNE plots labelled by cluster index. (C) TSNE plots labelled by major-categories. (D) TSNE plots labelled by sub-categories (The simulation data in B, C, D: 1. RDb.2; 2. RDb.3; 3. RDb.5; 4. RDb.6; 5. RDb.6.1). (E) The major-category classification accuracy curve. (F) The sub-category classification accuracy curve. sequencing depth.

The analytic results were visualized in TSNE plots, and the cells was marked with cluster index and true labels respectively (Figure 3B, 3C, 3D; Figure S1A, B, C). The classification accuracy for main category and sub category were calculated (Figure 3E, 3F). The major category accuracy quickly entered the plateau period, while the of sub-category accuracy is significant lower even the reads per cell reached 17324 and genes per cell reached 3906. However, the cluster number current scRNA studies is ∼40 sub-categories, while the average reads and genes detected were far less (reads per cell around 2000, genes per cell around 600). Accordingly, the depth of current platforms could effectively distinguish the main categories, while not for sub-categories.

### Parameters and analytic algorithms influence clustering accuracy in low depth

Since the sequencing depth current platforms is relatively unsatisfactory according to our previous results, the parameter and analytic algorithms used for analyses under low depth situation was evaluated. Firstly, simulated data was used for evaluation. Currently, the efficiency of specific-genes based annotation remains unknown. A simulation data with low accuracy (RDc.2.1, Table S5: Accuracy of major category: 0.6606445; Accuracy of sub-category: 0.2412109) which has a relatively acceptable cell cluster distribution (Figure 4A), was chosen for latter analysis. After differential gene analysis (see Materials and Methods), the top 20 specific genes of each cluster exhibited good discriminate performance (Figure 4B). However, only 85 genes out of the specific genes (716) overlapped with the 530 hemocyte-specific genes (Highly distinguishable genes for 42 sub-categories; Figure 4C), which meant that the cluster-specific genes only recovered less than a quarter of the true prior knowledge in ground truth (Figure 4D). CXCR1 and FCGR3B (Figure 4C), which are specific genes of neutrophils, were highly expressed in class 4 (Figure 4B), and the distribution of their expression was highly consistent with neutrophils (Figure 4E 1–3). CD2 (Figure 4C), a specific gene of T cell, was not identified while its expression was highly consistent with the T cells distribution (Figure 4E 4–5), which may result from the nonlinearity distribution of the T cell (Figure 4E 4). Thus, it is investigated that the type-specific genes of prior knowledge could describe the distribution of the type of cells in the scRNA-seq data in low depth condition. CD5 is highly expressed in three sub-categories of T cells: activated memory T cells, Tregs, and Teffs (Figure 4C). However, the distribution of the CD5 expression was not consistent with these cell types (Figure 4E 6–7). Therefore, the analyses showed that the results of scRNA-seq may not fully reproduce prior knowledge under low Subsequently, real data was used to estimate the impact of algorithms and parameters on clustering accuracy. We designed a sequencing scheme that five samples were sequenced twice after sequence libraries built (Table S1; see Materials and Methods) with different sequencing depth. The distance among the corresponding batches of each cell was quantified (see Materials and Methods). The clustering results (Figure S4) of identical cell varies (Figure 5A 3, 4): some of the identical cells in close proximity distributed on the contact line of two clusters and wrongly organized into two cluster, while some distributed in one cluster scope but still wrongly organized by analytic algorithm; There are some identical cells that are far apart but classified rightly in one cluster(Figure 5A 1, 2) while mis-classification is the main cases in the condition (Figure 5A 5, 6). The mis-classification rate of identical cells was quantified. The result showed that the mis-classification rates of some clusters were higher than 0.2 for all samples and even higher than 0.8 (Figure 5C). There is no biological bias between the two identical cells, thus the inconsistent rate is due to only sequencing and analytic processes. However, the cell individuals in the same cluster are heterogeneous, the mis-classification rate will be higher. The mis-classification rate showed a positive correlation with the cluster tightness (Figure 5B 1), while showed an opposite correlation with the distance from distribution centre to the cluster centre (Figure 5B 2). These patterns were also observed in simulation data. Two clusters, 4 and 8, which are away from the centre of the TSNE plot, were highly coincident with neutrophils cells and endothelial cells, respectively (Figure 5D 1, 5, 7). The cluster 7 near the centre was composed of three types of cells (Figure 5D 1–4). The cluster 9 which composed of three types of cells (Figure 5D 1, 6, 8, 9), was away from the centre, but relatively scattered, Collectively, the tightness and distribution of cluster in TSNE plot can provide some information about classification accuracy.

**Figure 4.**
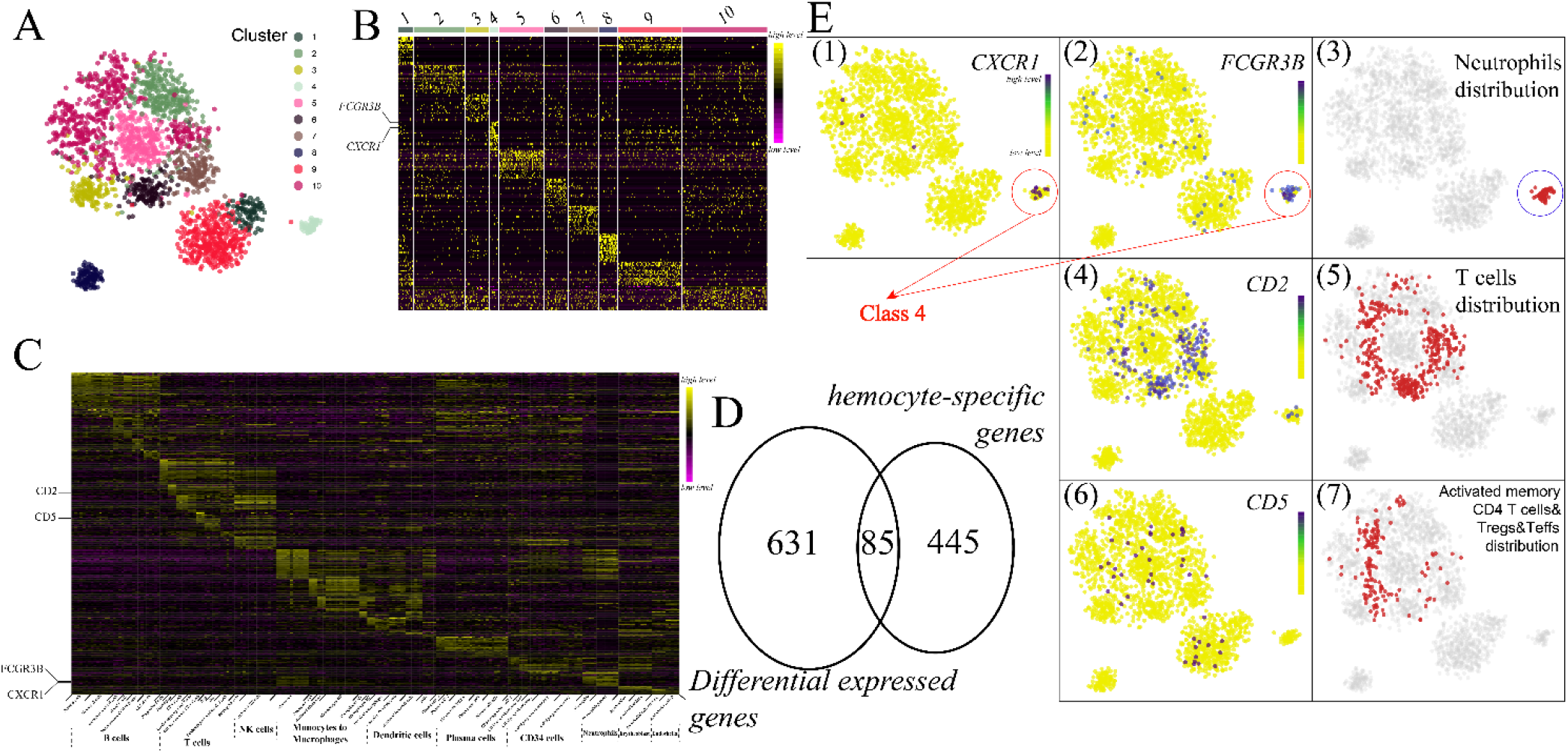
(A) TSNE plot of RDc.2.1 data (Table S5); (B) Heatmap of class specific-genes in RDc.2.1 data; (C) Heatmap of 530 hemocyte-specific genes in ground truth; (D) Venn diagram of class specific-genes and 530 hemocyte-specific genes; (E) Class, cell and gene expression distribution map in TSNE plots (1. CXCR1 gene expression abundance; 2. FCGR3B gene expression abundance; 3. Neutrophil cell; 4. CD2 gene expression abundance; 5. T cell; 6. CD5 gene expression abundance; 7. Specific T cell (activated memory T cells, Tregs, and Teffs))

**Figure 5.**
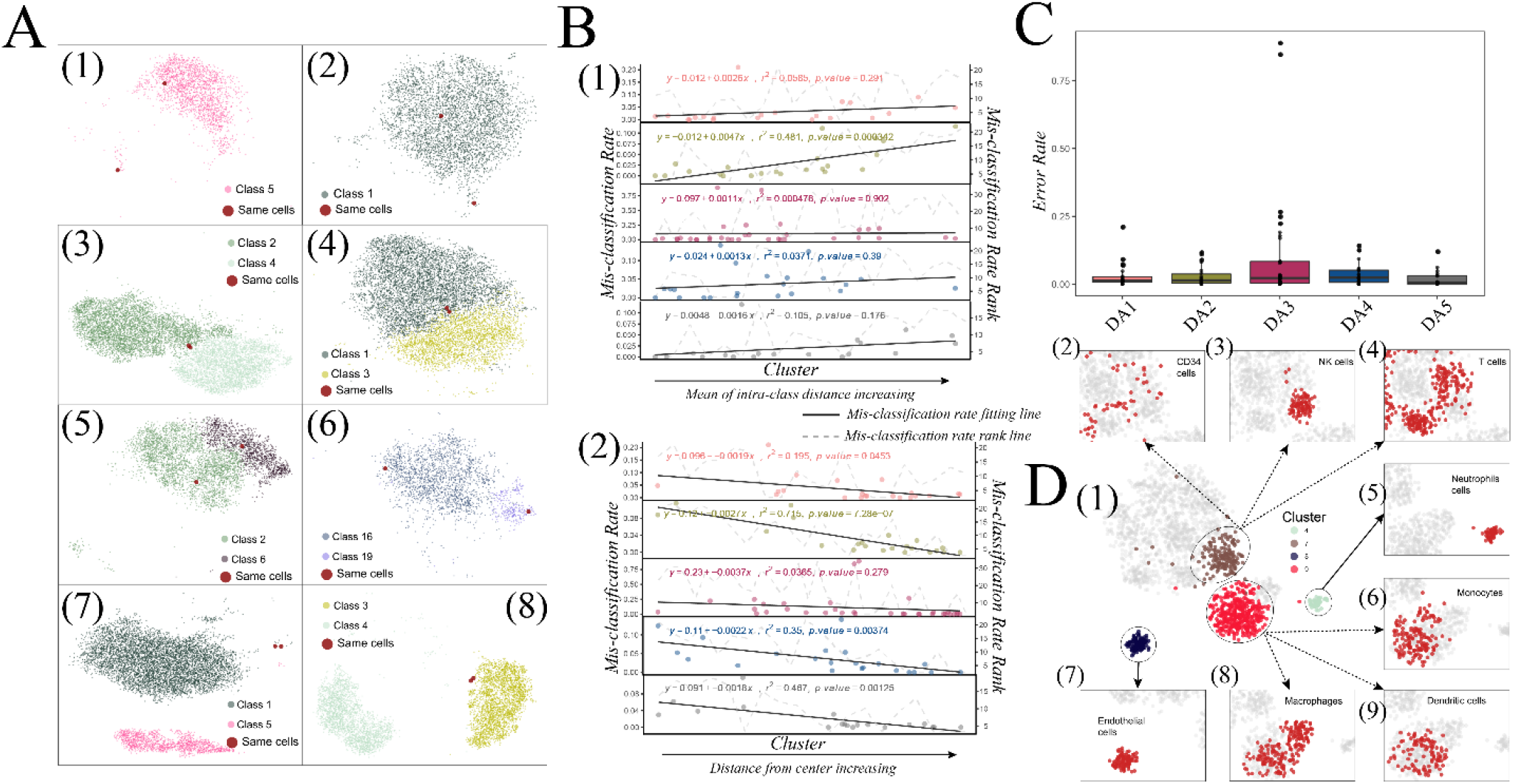
(A) The two identical cells on the TSNE plot. (The real data of each class: 1. DA2; 2. DA5; 3. DA2; 4. DA4; 5. DA1; 6. DA3; 7. DA1; 8. DA1) (B) Correlation plots of mis-classification rates with cluster-related parameters. (1. Correlation between mis-classification rate and mean distance between cells within clusters; 2. Correlation between mis-classification rate and distance of class centre to TSNE plot centre; Black line: fitted regression line; Dashed grey line: the rank of cluster-related parameter versus the rank of mis-classification rate) (C) Boxplot of the mis-classification rates of 5 samples. (D) TSNE plot of RDc.2.1 data (1. Ladled with 4, 7, 8, 9 cluster index; 2. CD34 cells; 3. NK cells; 4. T cells; 5. Neutrophil cells; 6. Monocyte cells; 7. Endothelial cells; 8. Macrophage cells; 9. Dendritic cells)

### Normalization algorithms that influence clustering accuracy in low depth

Similar to the real scRNA-seq data^[20]^, dropout values (count is 0) and low count (count is 1 or 2) values also consist of major part of expression matrix in simulation data. The dropout and low count ratios oppositely correlated with sequencing depths (Figure 6A, B). Consistently, more depth resulted in less drop-out and low count values in the real data (Table S1). A simulated dataset with similar depth was generated for latter analyses (Table S5). the data was normalized with 12 methods, followed by the identical dimension reduction and clustering methods (see Materials and Methods). It is observed that the different normalization algorithms had a considerable impact on clustering accuracy (Figure 6C, D). The TPM and edgeR^[30, 31]^, which were recommended in bulk-RNA seq analysis, performs worst among the algorithms. A preferred algorithm, scran^[32,33]^, which normalize the sub-clusters separately, does not outperform the others. Interestingly, a simple normalization method, scale (z-score normalization), performed better for sub-clustering. Specifically, log transformation improves the accuracy of normalization algorithms. Since log transformation and scale normalization are both Gaussian standardization method, thus, Gaussian transformation is recommended.

**Figure 6.**
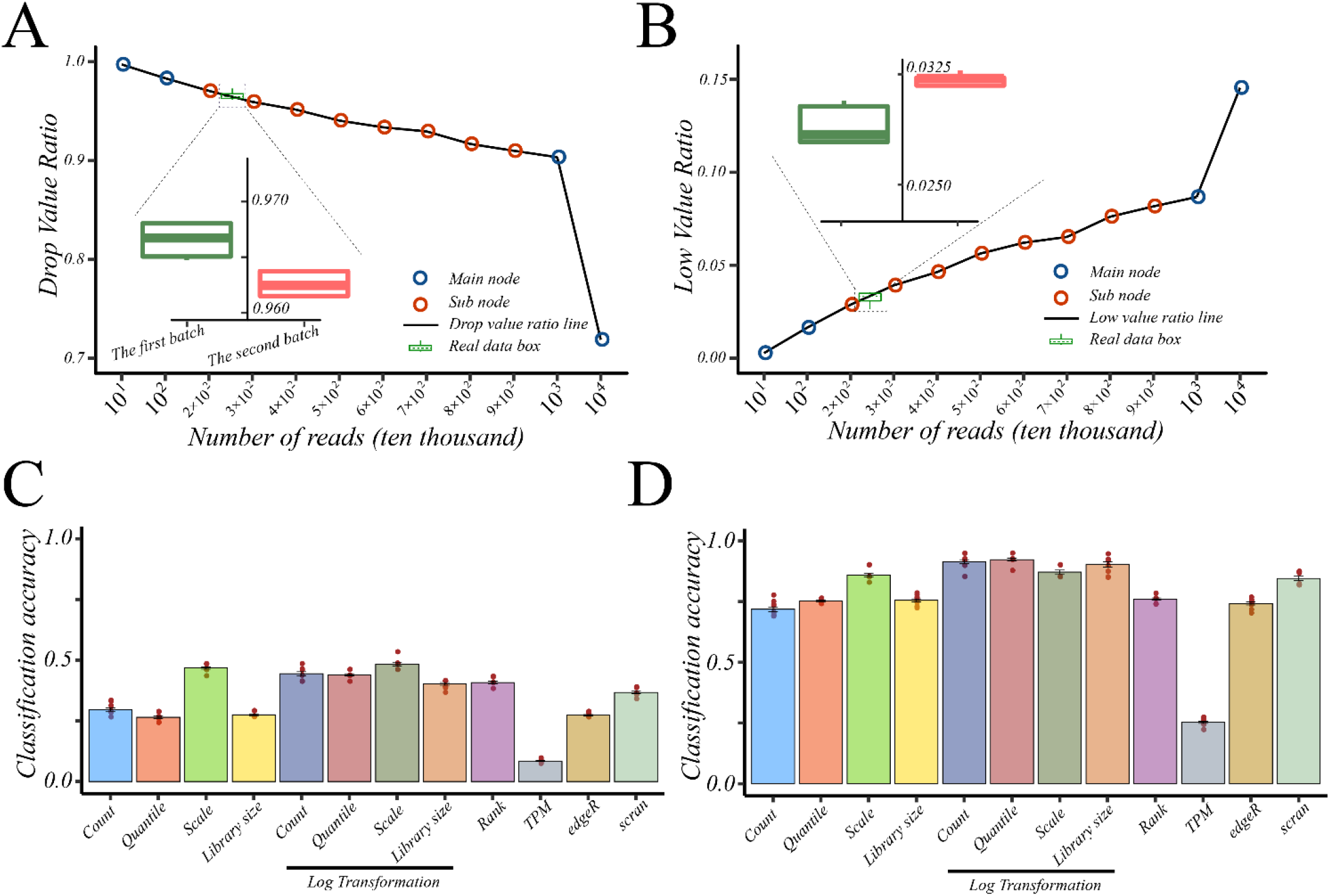
(A) Relationship between dropout ratio and number of reads. (B) Relationship between low value ratio and number of reads. (Blue point: main-gradient; red point: sub-gradient; boxplot: the two ratios of the two batches of real data) (C) Major category classification accuracy of 12 normalization methods. (D) Sub-category classification accuracy of 12 standardization methods.

### Dimension reduction and clustering algorithms that influence clustering accuracy in low depth

Currently, quite a few dimension reduction and clustering algorithms are available for researchers, while the performance of the methods is still obscure. The impact of 2 dimension reduction and 5 clustering algorithms (see Materials and Methods) were compared. To comprehensively investigate their performance, each normalization result of each dataset was enrolled in this step. As seen by the result (Figure 7A, Figure S3), som and hierarchical clustering is the least effective. The result of density clustering^[34]^ algorithm that utilizing TSNE coordinate as feature shows high accuracy in major-categories differentiation, while does not perform well in sub-categories differentiation, which indicated that in TSNE diagrams, sub-categories are often not effectively distinguished. The classification accuracy result of K-means algorithm exhibits steady regardless of any features employed for clustering. The result of Kmeans&Louvain algorithm performed relatively satisfactorily when using reduced component feature for clustering, while performances worse without dimension reduction (with all gene as feature). Combination of normalization and clustering algorithms significantly impacts the overall clustering accuracy. For instance, the kmeans&louvain algorithm performed better with scaled data (Figure 7C 1), while the kmeans algorithm performed better with quantile normalized data ((Figure 7C 2–3). The variance of classification accuracy also varies among clustering algorithm (Figure 7B, left panel: with different clustering feature; right panel: with different normalization method). Consistent with the previous result, K-means&Louvain is most unstable with different features for clustering, while K-means is the most stable one. In summary, the performance of the 5 clustering algorithms was somewhat divergent, while K-means&louvain and K-means should be the best choice in current workflow, and K-means is a more prudent option.

**Figure 7.**
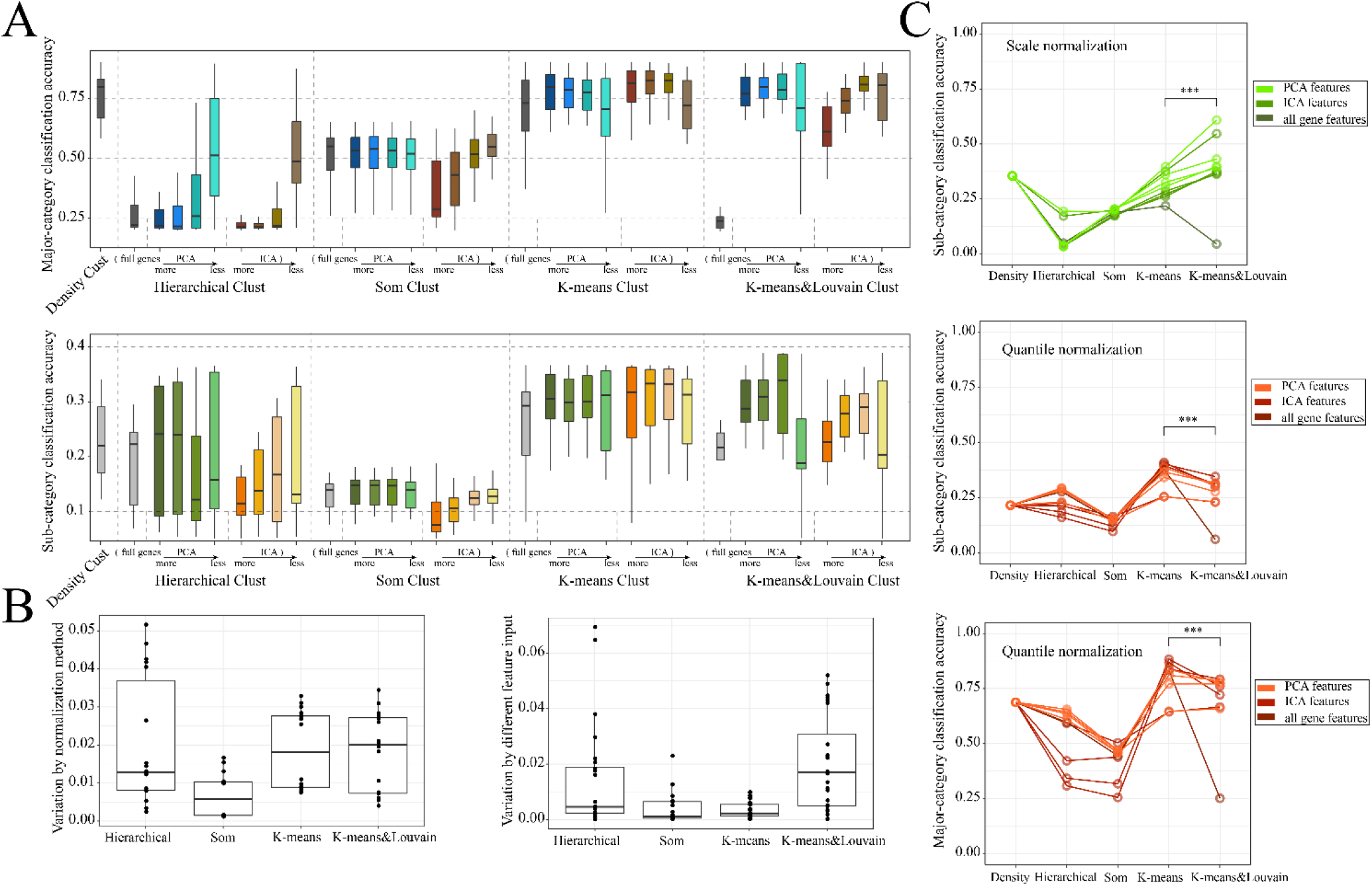
(A) The boxplot of classification accuracy with different combination of dimension reduction method and clustering method (upper panel: major-category classification accuracy; lower panel: sub-category classification accuracy; the feature numbers labelled with ‘more to less’: 100, 70, 40, 10). (B) The boxplot of classification variance with different clustering method (left panel: variance calculated by different normalization method; right panel: variance calculated by different dimension reduction method). (C) The line plot of classification accuracy of different clustering methods (upper panel: Sub-category classification accuracy of scale normalized data with different clustering method; middle panel: Sub-category classification accuracy of quantile normalized data with different clustering method; lower panel: major-category classification accuracy of quantile normalized data with different clustering method).

## DISCUSSION

This project proposed a program (SSCRNA), which utilizes a pre-defined ground truth to simulates the scRNA-seq raw data. SSCRNA mimics the real sequencing process at all stages, allowing for generating raw data according to the parameters of different sequencing platform. The comparison of simulation data with real data verified the reliability of the program. A ground truth which obtained by augmenting collected data was employed for simulation. We used the simulated data to examine the effect of sequencing depth and analysis workflow on classification accuracy. The test result of sequencing depth suggests that the real data (10,000 cells) need at least 50 million reads to achieve better classification results (the classification accuracy of 7 major categories is close to 1, and of 42 sub-categories is more than 0.5). The test result of analysis workflow suggests that gaussian normalization is suitable for current workflow and K-means clustering is more stable than K-means&Louvain clustering. The scope of the conclusion is limited to the cluster-annotation way. For some other annotation methods that may emerge in the future, it is unknown which normalization algorithm will perform better. Because it is believed that a smaller deformation for the raw data which retain more information might enable a higher potential for upper limits on classification accuracy.

For the fitting of single-cell data properties, researchers have developed several simulation algorithms, including splatter, SPsimSeq, SPARSim and SymSim, which are all count simulators^[14–17]^. The splat algorithm is recommended in splatter package which also inherits a variety of simple algorithm, such as lun2, scDD, etc^[35–38]^. The splat algorithm assumes that the gene expression profile is based on a negative binomial distribution, and estimates outlier probability, library size and dropout indicator from real data to generate the observed count as simulation data. SimSeq is non-parameter method, which make simulation by sampling from real data. Based on this, SPsimSeq was aroused as a semi-parameter method, which make use of Gaussian-copulas to retain the between-genes correlation structure. The SymSim method assumes that the individual gene’s expression follows the stationary distribution of the two-state kinetic model, which using three parameters: expression on, off probability and transcription rate, and specifies the cell state by EVF (extrinsic variability factors). SPARSim, on the other hand, constructed a single-cell count matrix model with a gamma-multifactor hypergeometric distribution model. The common denominator of these method is that they assume single-cell expression data satisfy numerous statistical models, and estimate the probability distribution of genes through several parameters from real data, and randomly sample from this distribution to generate simulation data.

However, for the data with abundant mixed types which may distribute differently, the algorithm with a relatively simple statistical model and small number of parameters is unlikely to simulate accurately. First, Due to the lack of single population expression data, these algorithms cannot accurately estimate model parameters. In our project, single population expression data were collected in large quantities and single genes were amplified individually, significantly maintaining the properties of single cell expression data. Second, the evaluation of the previous method is limited to the comparison of the parameters estimated from the overall distribution and lacks the interpretation of cell population and individual characteristics. Here is an extreme example being the random swapping of data positions in the count matrix, which will not change the distribution of various parameters (sparsity, coefficient of variation, etc.), but the matrix after the swap is not consistent with the original matrix. Moreover, although the data processing of single-cell sequencing is more consistent with bulk-sequencing data, many parameters, such as sequencing error, mapping efficiency and sequencing depth, may affect the analysis results in different level. The previous algorithm ignores the mapping of count value to reads, while SSCRNA program allows a more complete integration of the whole process.

The quantification of the “true state” of the cell population used in the construction of ground truth were derived from the dataset of array platform. There was a certain degree of subjectivity, such as quantitative relationship between the signal intensity of the probe and the true mRNA number. The diversity of gene sequences also introduces noises in the actual sequencing process, which causes lower mapping accuracy. But this bias was not implemented in SSCRNA simulator. This study mainly discusses sequencing depth in scRNA-seq analysis, which should be the parameter that has the most obvious impact on accuracy. Other parameters in the sequencing and analysis process need further exploration, such as the error propensity of different sequencing platforms, different cell barcode estimation algorithms and the types of errors that may be introduced by different library building processes.

Several potential directions of analysis can be pursued in further analysis, such as exploring the patterns of single-cell expression, screening for new methods for single-cell analysis and testing the effectiveness of differentiation-related algorithms^[39, 40]^. It is necessary to screen for more relevant algorithms that may generate better results, and SSCRNA program makes the screening process possible.

## MATERIAL AND METHODS

### SSCRNA program

SSCRNA program is implemented using python, and utilizes the human-defined ground truth (Figure 1B 1) to perform scRNA-seq simulation process which consists of two parts. The first part is library building simulation (Figure 1B 2); the second part is sequencing process simulation (Figure 1B 3–5).

In the first part, the final constructed sequencing library is made up of four sub-libraries: the fragment library, PCR control library, UMI library, and cell barcode library (Figure 1B 2). The construction of sequencing library is based on human-defined ground truth (an expression matrix containing mRNA numbers is defined artificially according to some rules or assumptions of actual single cell expression profiles; Figure 1B 1). The human reference genome GRCH38 (gencode.v32 transcripts) is used as reference sequence for the genes in ground truth. SSCRNA program allows a certain size of deletion at the head and tail of the genes (may be introduced during the real process of cell lysis and reverse transcription; default set to 10bp). A single fragment file is composed of all the gene fragments (sequences after random depletion; Figure 1B 2 upper left panel) of a single cell, that is, the first column is genes name, and the fragments are recorded (as bed format) behind the genes. The fragment files of all cells constitute the fragment library. And for each cell, we set a cell barcode of specific length (default is 25bp, and any two barcodes are at least two base units apart) to form the cell barcode library. Then for each sequence in fragment library, we randomly assign a UMI (unique molecular identifier) of a set length (default is 8bp) to form a UMI library, which has the same structure as the fragment library (Figure 1B 2 lower right panel). After the above three libraries have been built, the PCR amplification simulation is performed, to make up the PCR control library (Figure 1B 2 lower left panel). PCR control library also has the same structure as the fragment library, of which the number of fragments and the information of random mutation (the position, type and outcome of the mutation) are marked behind each gene (the default amplification number is 8 (3 cycles amplification) and no mutation occur). At this point, the construction of the simulation sequencing library composed of four sub-libraries is completed, the size of the simulation sequencing-library is obtained by summing the number of all fragments in PCR control library.

The second part was the on-machine sequencing process simulation (Figure 1B 3–5). In the sequence selection process, fragments (each fragment in PCR control library was encoded) are randomly selected, and the corresponding UMIs and cell barcodes are extracted from the sub-libraries. According to the combination rule of sequence elements (adaptor, UMI, cell barcode, gene sequence), full-length sequence is generated (Figure 1B 3). In read generating process, we introduce the probability model of reading errors, which is stored in error control file (Figure 1B 4 left panel). The assumption is that the probability of an error is decided by three conditions: the type of base, its position in the sequence, and the type of error (addition; deletion; substitution). The quality control profile was also introduced (Figure 1B 4 right panel) to control the base quality in reading process. The assumption is that the expectation of read quality is decided by three conditions: base type, its position in the sequence, and whether or not a read error occurs. Read generating simulation is proceeded from both ends of the full-length sequence, and is constrained by error and quality control file. After generating a sequence of a certain length (default 150bp), the reading process for each full-sequence is aborted, and the sequence and the quality information were imported into the raw data files (Figure 1B 5).

### Construction of ground truth

From the GPL96 platform, we collected 225 samples from 11 datasets (Table S2), and the samples were classified into 11 major-categories and 42 sub-categories according to the type of enriched cells. Each sub-category (the sample size is at least 3) was normally amplified to 50 samples based on the mean and variance of the included genes. Thus, finally a dataset with 2100 cells was generated.

### The parameters in single-cell sequencing simulation

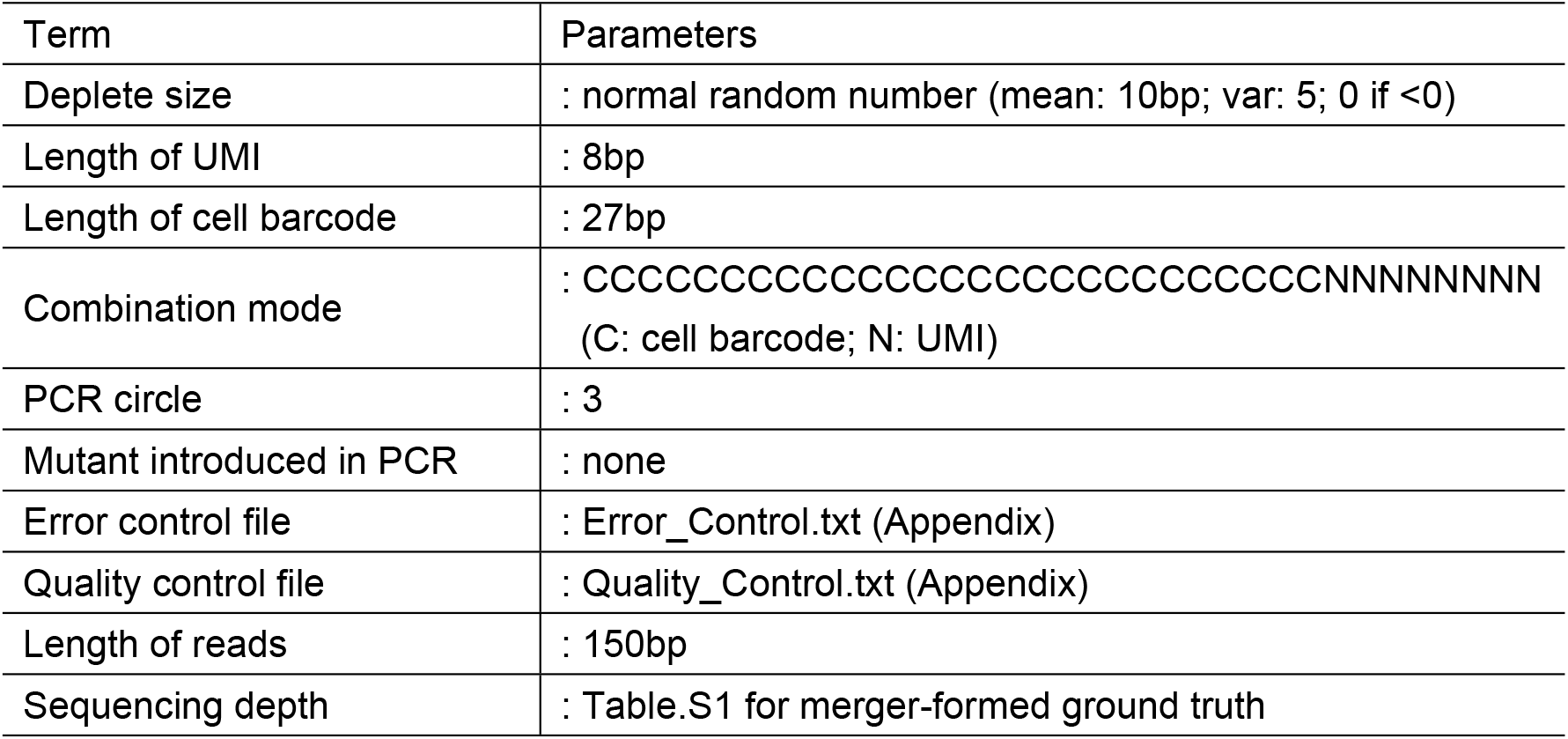
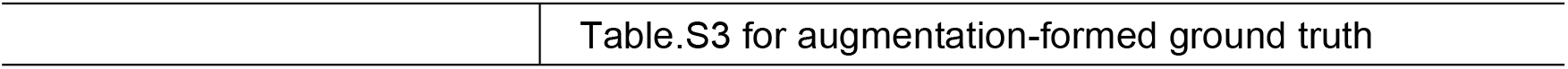

### Real data collecting

Thanks for School of Basic Medical Science, Fudan University for providing 5 sequencing libraries of human liver tissue samples which was constructed using 10 × genomics platform. Each sequencing library was then sequenced twice on the Illumina platform and the second batch was required to have two times size than the first batch (Table S1).

### Correlation calculation between samples of collected dataset on GPL96 platform

Whole genes and 530 hemocyte-specific^[29]^ genes were used to calculate the correlation between collected samples (Table S4). The cor function in R environment was used to calculate Pearson correlation coefficient between samples, and the data was scaled before correlation calculation.

### Differential analysis for class-specific genes

Limma package^[41]^ was used to calculate differential expression gene for each class in one-VS-others way. If the gene is obtained as differential gene for more than four clusters, the gene is deleted, and the remaining genes were viewed as the class-specific genes.

### Default procedure for single cell analysis Raw data processing processes

the putative cell barcode was estimated by whitelist function in UMItools, and the cell barcode and UMI were extracted using extract function. STAR software was used to map the reads to reference genome (GRCH38). The featureCounts software was used determine the gene number according to the map results (gencode.v29). the count function in UMItools was used to eliminate the polarization effect during amplification process and to obtain the final scRNA-seq expression sparse matrix.

### Count data processing processes (Default workflow)

R language was used for the subsequent analysis of the expression matrix. The library.size.normalize function of the phateR package^[42]^ was used to make global library size normalization. The prcomp function was used to reduce feature dimension and the top 30 feature vectors was selected. The cells were clustered using the Rphenograph package^[43]^, which was based on the k-means&louvain algorithm. TSNE plots^[44]^ were used to display the distribution of cells which was incorporate in the Rtsne package.

### Standardization, dimension reduction and clustering methods

All the algorithms were implemented in R environment. The following is explanation of the function for each algorithm.

### 12 standardization methods

Count data: Expression matrix obtained by the count function in UMItools Quantile: normalize.quantiles function in preprocessCore packages

Scale: scale function

Library size standardization: library.size.normalize function of the phateR package

Log transformation: log10 function

Rank standardization:rank function

TPM standardization: the formula of count to TPM (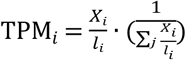; | represents transcript length; i represents gene number; j represents cell number) was used to obtain the TPM matrix.

edgeR standardization^[31]^: using each cell as a sample, standardized factors was calculated using the calcNormFactors function in the edgeR package, common dispersion was calculated using estimateCommonDisp function, and intergenic range dispersion was calculated using estimateTagwiseDisp function. The estimated pseudo counts matrix were multiplied with the standardized factors to obtain the final standardized data.

scran standardization^[33]^: The SingleCellExperiment function in scran package was used to convert expression matrix to SingleCellExperiment objects, and the quickCluster function was used to sub-cluster cells. Then the computeSumFactors function was used to calculate standardized factors within each subclasses, and finally, the normalize function was used to complete the standardization.

### 2 dimension-reduction methods

PCA: prcomp function

ICA (Independent Component Analysis)^[45]^: fastICA function in fastICA package

### 5 clustering methods

Density cluster: findClusters function in densityClust package^[46]^

Hierarchical cluster: hclust function

Som (self-organized map)^[47]^ cluster: som function in som package

K-means cluster: kmeans function

K-means&louvain cluster: Rphenograph function in Rphenograph package

## Supplementary figure

**Figure S1.**
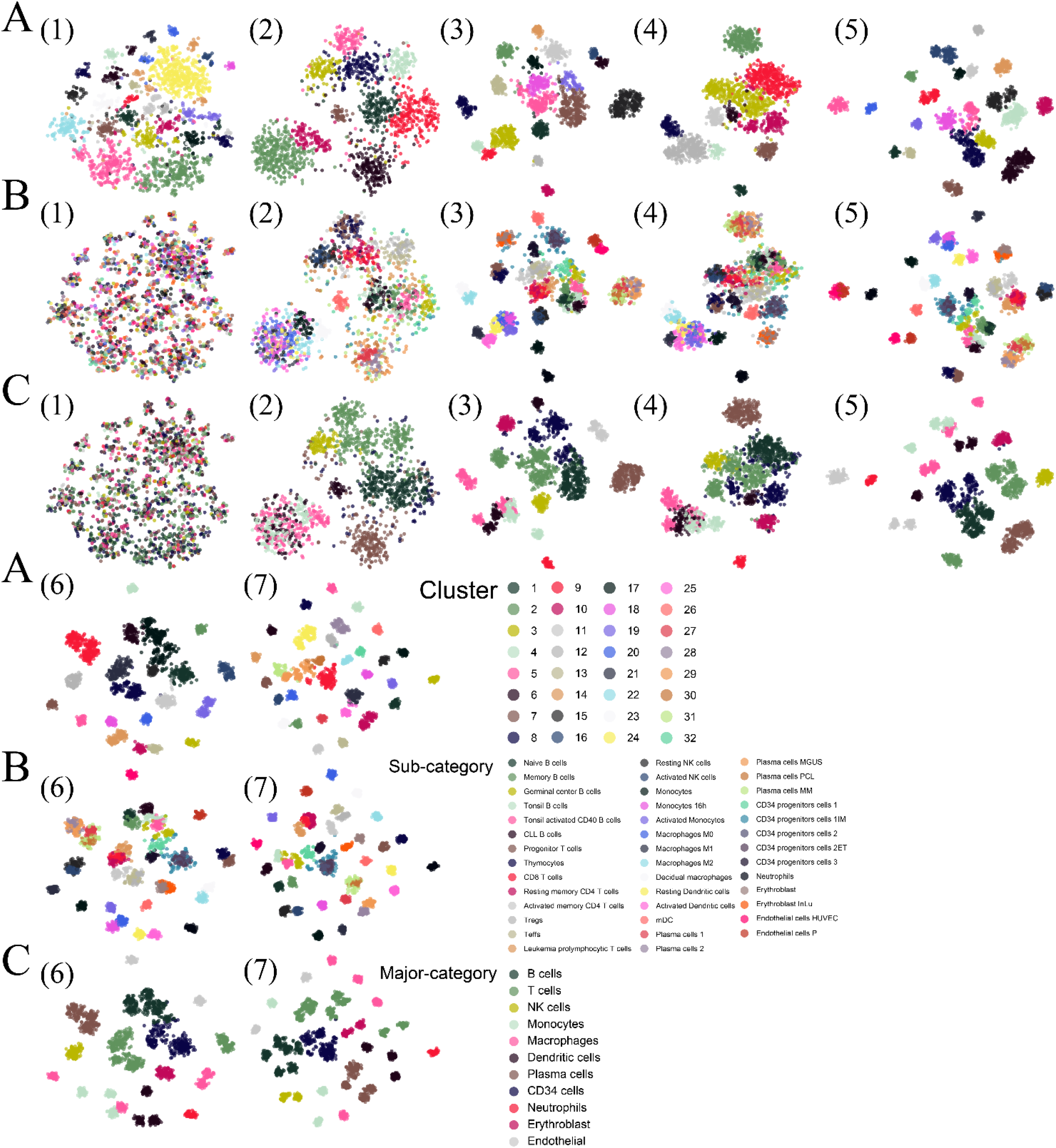
(A) TSNE plots labelled by cluster index. (B) TSNE plots labelled by major-category labels. (C) TSNE plots labelled by sub-category labels. (Simulation data for A, B, C: RDc.1; 2. RDc.2.2; 3. RDc.2.4; 4. RDc.2.6; 5. RDc.2.7;6. RDc.2.8;7. RDc.4)

**Figure S2.**
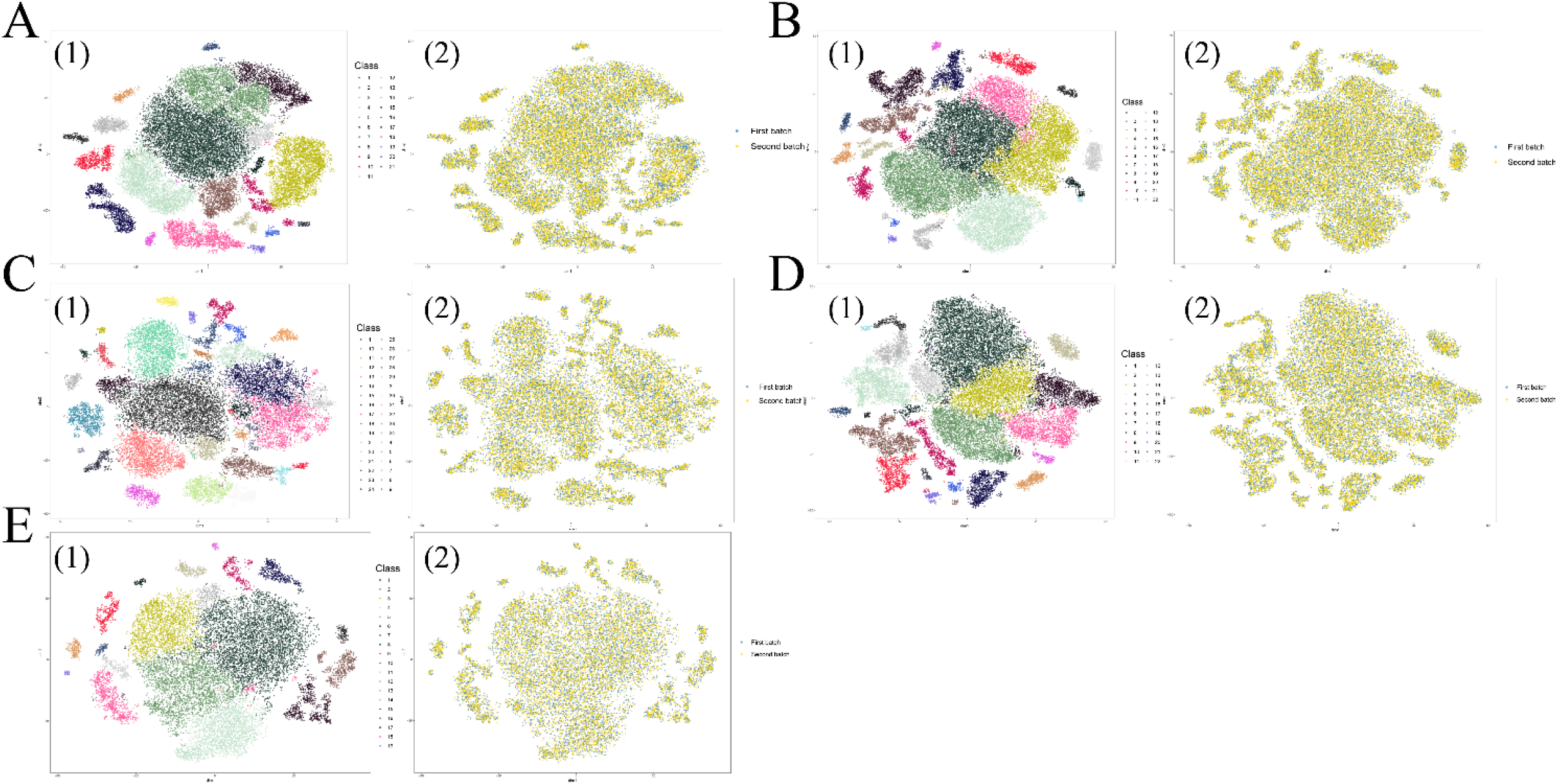
(A) TSNE plots of cell distribution of DA1. (B) TSNE plots of cell distribution of DA2. (C) TSNE plots of cell distribution of DA3. (D) TSNE plots of cell distribution of DA4. (E) TSNE plots of cell distribution of DA5. (1. Cells labelled by class labels; 2. Cells labelled by sequencing batch)

**Figure S3.**
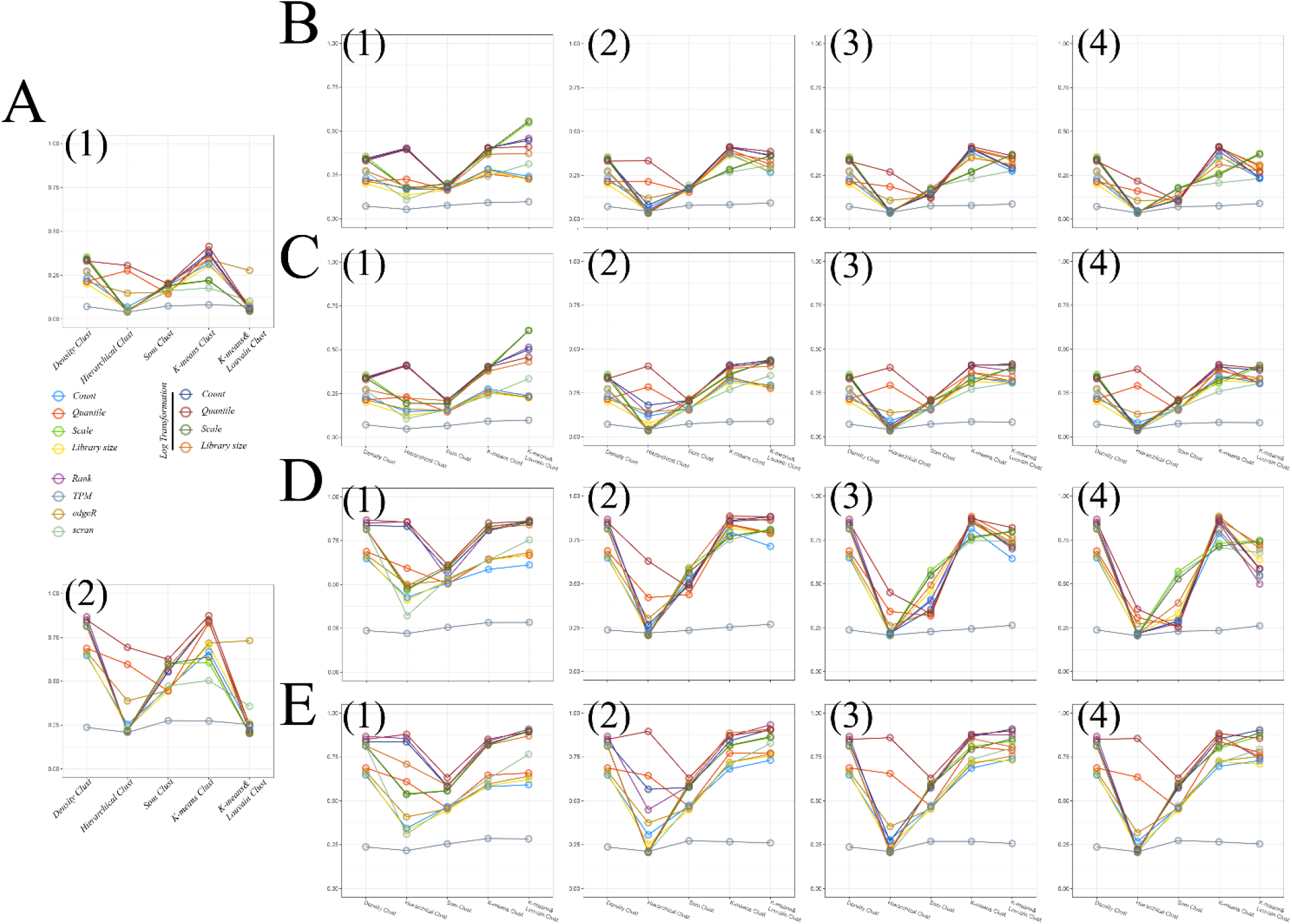
(A) The classification accuracy of different clustering methods without dimension reduction. (The colours correspond to the different normalization methods; 1. Accuracy for sub-category; 2. accuracy for major-category) (B) The sub-category classification accuracy of different clustering methods with ICA feature. (1. 10 features; 2. 40 features; 3. 70 features; 4. 100 features) (C) The sub-category classification accuracy of different clustering methods with PCA feature. (1. 10 features; 2. 40 features; 3. 70 features; 4. 100 features) (D) The major-category classification accuracy of different clustering methods with ICA features. (1. 10 features; 2. 40 features; 3. 70 features; 4. 100 features) (E) The major-category classification accuracy of different clustering methods with PCA feature. (1. 10 features; 2. 40 features; 3. 70 features; 4. 100 features)

## Availability and Implementation

SSCRNA: https://github.com/liuyunho/SSCRNA-v1.0

## FUNDING

This work was supported by:

the National Natural Science Foundation of China (#91846302),

the National Key Research and Development Program of China (No. 2016YFC0901900),

the Shanghai Municipal Commission of Health and Family Planning,China(Grant No. 2018ZHYL0104).

## DATA AVAILABILITY

Data (Table S1) has been uploaded to GEO.

Reviewer link: https://dataview.ncbi.nlm.nih.gov/object/PRJNA681982?reviewer=odr2qng06esoi9v1gtdacctsj4

